# Rapid colour changes in a tiny threatened gecko do not impede computer-assisted individual recognition

**DOI:** 10.1101/2022.03.16.484634

**Authors:** Cindy Monnet, Théo Dokhelar, Julien Renet

**Author notes:** Correspondence: Julien Renet, Conservatoire d’espaces naturels de Provence-Alpes-Côte d’Azur, Pôle Biodiversité régionale, 18 avenue du Gand, 04200 Sisteron, France.

## Abstract

Photo-identification is a non-invasive method used for recognising wild animals with distinctive and stable patterns over time. This method is now widely used for capture-recapture wildlife monitoring. However, in species exhibiting rapid colouration changes, the evolving body patterns can lead to errors in individual recognition. In this study, we assessed the effect of dorsal physiological colour change of the tiny threatened European leaf-toed gecko (*Euleptes europaea*) on the performance of Wild-ID and Hotspotter, the two most commonly used individual recognition software for wildlife monitoring. We exposed 30 European leaf-toed geckos to several semi-controlled parameters (substrate type, temperature and light from natural diurnal/nocturnal cycles) in order to characterise the extent of variation in dorsal colouration, by standardised reflectance measurements. The colour of the substrate had a significant effect on individual reflectance changes. Body temperature also seemed to significantly affect the reflectance but the experimental conditions did not allow us to clearly distinguish the effect of temperature and light. For each of the 30 geckos, four photographic databases (*n* = 4*280) were then analysed by both software packages, under two extreme reflectance conditions. Despite the large changes in individual reflectance, Wild-ID and Hotspotter proved to be extremely reliable with a 100% recognition rate. The analysis of similarity scores suggests that Hotspotter is less sensitive to chromatic variation than Wild-ID. We provide here the first evidence that physiological colour change is not a barrier to computer-assisted individual recognition. This study advocates the use of Hotspotter for monitoring populations of European leaf-toed geckos and other saurians that generate significant colouration change over a short time.

## Introduction

Photo-identification is a non-invasive and low-cost method widely used in wildlife capture-recapture monitoring (Jackson *et al*., 2006; Bengsen, Butler & Masters, 2011; Başkale & Kaya, 2012; Pace, Corkeron & Kraus, 2017; Schofield *et al*., 2020). It limits the pain and discomfort caused by more invasive individual marking techniques (e.g. toe clipping) to only the capture stress induced by photography. This method however requires distinctive body patterns (e.g. spots, stripes, notches) from the studied species that remain stable over time (Bolger *et al*., 2012). Although visual matching of photos is considered a valid methodology to monitor small populations during limited time periods, it can become tedious, time consuming and prone to misidentification when used to monitor large populations over several years (Morrison *et al*., 2011; Cruickshank & Schmidt, 2017).

In order to increase efficiency and reliability, several pattern recognition algorithms have been developed and integrated into photo-identification software. Such software allows users to automate the matching between two images of the same individual based on characteristic body patterns or easily identifiable morphological features (e.g. scale morphology) (Sacchi *et al*., 2016). Wild-ID (Bolger *et al*., 2012) and Hotspotter (Crall *et al*., 2013) are two of the most commonly used open source individual recognition software for the study of animal population dynamics.

Based on the Scale-Invariant Feature Transform (SIFT) algorithm, Wild-ID has been successfully used in a wide array of taxa (amphibians, reptiles, mammals) (Bolger *et al*., 2012; Cross *et al*., 2014; Morrison & Bolger, 2014; Matthé *et al*., 2017; Bardier *et al*., 2020), even for species exhibiting complex body patterns (Bendik *et al*., 2013; Renet *et al*., 2019). Hotspotter was developed from the SIFT and Local Naive Bayes Nearest Neighbor (LNBNN) algorithms. It has also shown high reliability in fishes (Crall *et al*., 2013), mollusks (Barord *et al*., 2014), insects (Quinby, Creighton & Flaherty, 2021), reptiles (Dunbar *et al*., 2021; Tabuki *et al*., 2021) and mammals (Crall *et al*., 2013; Lea *et al*., 2018; Nipko, Holcombe & Kelly, 2020).

Not all individually identifiable patterns, however, are stable over time. Biologists tend to avoid visual or computer-assisted matching photo-identification for individual recognition when dealing with species exhibiting rapid colouration changes in response to environmental stimuli, such as temperature (Langkilde & Boronow, 2012; Smith *et al*., 2016), stress (Boyer & Swierk, 2017; Lewis *et al*., 2017) and social interactions (Stuart-Fox & Moussalli 2008; Ligon & McGraw, 2013; Ligon, 2014). This physiological colour change is induced by the movement (aggregation or dispersion) of pigment granules in the chromatophores in a fast process that can take only a few seconds (Bagnara & Hadley, 1973). This phenomenon is widespread in the animal kingdom, including arachnids (Wunderlin & Kropf, 2013), insects (Sun *et al*., 2017), crustaceans (Kronstadt, Darnell & Munguia, 2013), cephalopods (Ikeda, 2021), fishes (Sköld *et al*., 2016), amphibians (Sköld, Aspengren & Wallin, 2013) and reptiles (Hadley & Goldman, 1969). Although the use of photo-identification for such species would increase individual recognition error rates and bias capture-recapture based demographic studies, to our knowledge, no studies have quantified the impact of chromatic variation on the reliability of individual recognition software.

Two types of errors are known in photo-identification. FARs (False Acceptance Rates = false positive errors) is the frequency of falsely matching two images of different individuals. FRRs (False Rejection Rates = false negative errors) is the frequency of failing to recognise two images of the same individual (Jain, 2007, Morrison *et al*., 2011; Bolger *et al*., 2012; Bendik *et al*., 2013). The most suitable photo-identification software for monitoring a species can therefore be determined by assessing such error rates (Sacchi *et al*., 2016).

The main objective of this study is to assess the impact of chromatic variation on the reliability of two popular individual recognition software (Wild-ID and Hotspotter). To achieve this, we characterised the degree of variation in the dorsal colouration of the European leaf-toed gecko (*Euleptes europaea*), a small endangered gecko endemic to the western Mediterranean. A sub-objective was also to understand the factors causing chromatic variation in this species. We finally provide methodological recommendations for the individual monitoring of European leaf-toed geckos and other saurians that can rapidly colour change.

## Materials and methods

### Study species

The European leaf-toed gecko (*Euleptes europaea*) is the smallest European gecko (snout-vent length < 5cm) and is endemic to the western Mediterranean Sea (Delaugerre, 1997). This gecko is a poikilothermic and strictly nocturnal species mainly inhabiting crevices and narrow faults of rocky environments (Delaugerre, 1981a). Nevertheless, this species seems to leave the rocky substrate to disperse into the vegetation during the warmest nights (Delaugerre, 1992). Melanophores (chromatophore cells that store melanin) are mainly responsible for the pigmentation of the European leaf-toed gecko (Delaugerre, 1981b). The dorsal side, with cream to black pigmentation, has a constellation of lighter scales reminiscent of the Milky Way galaxy. A pale dorsal line with light transverse bands is usually found on the back of individuals. This species has the particularity of rapid colour changes, from a light to a darker colouration, revealing or attenuating the dorsal patterns.

On a global scale, the European leaf-toed gecko is classified as Near Threatened by the International Union for Conservation of Nature (IUCN) Red List (Corti *et al*., 2009). However, a significant proportion of populations located on the margins of the distribution range seems to have undergone demographic collapses (Delaugerre, Ouni & Nouira, 2011; Costa, Oneto & Salvidio, 2019). This is particularly the case in south-eastern France, where populations are exposed to threats that are mostly anthropogenic (forest recolonisation, habitat destruction and fragmentation, urbanisation, introduction of co-occurring species, etc.) (Delaugerre *et al*., 2011; Renet *et al*., 2013). As a result, the species is considered Endangered in the French region Provence-Alpes-Côte d’Azur (Marchand *et al*., 2017) and is currently the subject of a regional conservation strategy involving various managers of natural areas (e.g. the Calanques and Port-Cros National Parks and the French coastal protection agency).

At present, no population monitoring at the individual level has been carried out because no permanent marking techniques have been conclusive for this species. The species’ small size makes it impossible to use transponders and the moulting phenomenon and rubbing in crevices prevent the persistence of any colour markings.

### Experimental design

In order to study chromatic variation in the European leaf-toed gecko, 30 adults were captured in May 2021 on the Lérins archipelago (islet of La Tradelière, commune of Cannes, France). The individuals were kept in captivity for four days in six 35×27×30cm terrariums installed out of direct sunlight exposition (Supporting Information Figure S1). Each terrarium contained a group of five individuals (males and females combined). These groups were kept together for the whole duration of the experiment. During captivity, the individuals were fed with invertebrates collected on the islet of La Tradelière.

To maximise the chances of obtaining a wide range of colouration, individuals were exposed to two substrate types subject to diurnal and nocturnal cycles implying natural light and temperature variations. The first three terrariums were composed of four walls coated with light-coloured cement with the bottom lined with mineral material from the islet La Tradelière. The other three terrariums were composed of four walls covered with topsoil and plant material was placed in the bottom of the boxes and collected on La Tradelière. All the terrariums were also equipped with two transparent Plexiglas walls perforated for aeration. This experimental design allowed us to mimic two micro-habitats used by the European leaf-toed gecko : a light-coloured rocky micro-habitat (RMH) and a dark-coloured vegetated micro-habitat (VMH) (Supporting Information Figure S1).

The substrate temperature and external body temperature of the individuals were measured with a FLUKE® 62 Max infrared thermometer before each manipulation (Supporting Information Tables S1 and S2). Individual recognition was based on a number inscribed on the ventral side of the individuals using an Edding® brand paint pen (non-toxic). Physiological colour change was assessed from photographs of the back of the individuals. Pictures were taken in a photographic studio offering standardised conditions (position of the animal, distance between the digital camera and the animal, angle of view, luminosity, etc.) (Supporting Information Figure S2). A first handler held the gecko while a second one took the photograph of the dorsal side. The photos were taken in RAW format (dimensions 4040 × 3016) in macro mode using an Olympus® TG-6 camera equipped with an LG-1 LED ring. The experiment took place over four consecutive nights and days. The different groups of individuals changed terrariums to ensure similar exposure to all investigated conditions, following the protocol described in Supporting Information Figure S1. Individual survival during the experiment was 100%.

### Reflectance analysis

Reflectance measurements are used to characterise the colours of a species’ patterns (Endler, 1990). By comparing such colouration to a standard greyscale, it is possible to measure an animal’s darkness (Hamilton, 2005). Total reflectance is the percentage of white light reflected from a surface (Hamilton, Gaalema & Sullivan, 2008). Using a photograph, the reflectance of each pixel is measured, ranging from 0 (black) to 1 (white) and corresponding to a reflectance of 0 to 100%. The average reflectance is then defined as the sum of the reflectance values of all pixels included in the selection divided by the number of pixels. In order to quantify the change in physiological colouration of the 30 geckos, the average dorsal reflectance of each individual was determined from photographs of the night 1 (N1) to day 4 (D4) (excluding day 1) i.e. under different light conditions (day/night), temperatures and substrates. In this way, the pairs of images exhibiting the maximum (max) and minimum (min) reflectance differences were identified for each individual.

All photographs were converted to Tagged Image File Format (TIFF) using XnConvert 1.90.0 (XnSoft Corp.) and then standardised to 8-bit greyscale in ImageJ 1.53a (Schneider, Rasband & Eliceiri, 2012). Images were then calibrated according to the method proposed by Hamilton *et al*. (2008) using the standard Tiffen Q-13 greyscale positioned next to each individual, with 20 shades representing grey reflectance values equal to 0.891 (89.1%) down to 0.011 (1.1%) (Supporting Information Figure S2). The average reflectance of the animal’s dorsal side was obtained by delineating the contours of the back, from the insertion of the front legs to the insertion of the back legs. The reflectance of the walls of the RMH and VMH terrariums corresponded to 0.386 and 0.071 respectively.

### Study of chromatic variation

We used generalised linear mixed models (GLMMs) to determine the effect of the substrate of the terrariums used, the body temperature of the individuals and the interaction between these two variables on the reflectance of our individuals. The collinearity of the different collected variables (substrate type, body temperature, substrate temperature, light) was analysed and variables with correlation coefficients greater than 0.7 were excluded from the analyses to avoid inaccurate estimation of coefficients in models. Thus, the substrate temperature and light variables were removed because they were both highly correlated with body temperature values.

Substrate type (*n* = 2), body temperature (*n* = 149) and the interaction of these variables were treated as fixed effects, while the individuals and the terrarium units were treated as random effects. Reflectance measurements (*n* = 149) were used as the sampling unit. GLMMs were based on data from day 2 (D2) to day 4 (D4) (i.e. two nights and three days) of the experiment and fitted using the glmmTMB function in the R package glmmTMB (Magnusson *et al*., 2017). The global models were fitted using a Gaussian distribution. Using the dredge function in the MuMIn package (Barton, 2020), a set of models was constructed representing all combinations of explanatory variables based on the global models. All models with an AIC delta of less than two relative to the best-fit model were considered to have substantial statistical support and are therefore considered for interpretation (Burnham & Anderson, 2002). The relative importance of each explanatory variable was assessed using the sum of Akaike weights (Σwi), showing the contribution of each predictor to the construction of candidate models (Johnson, 2000; Tonidandel & LeBreton, 2010). To check the quality of fit of our selected models, we calculated the marginal R^2^ (Rm^2^), estimating the variance explained by the fixed factors, and the conditional R^2^ (Rc^2^), estimating the variance explained by both the fixed and random factors (Nakagawa, Johnson & Schielzeth, 2017), using the performance package (Lüdecke *et al*., 2021). Pairwise comparisons of EMMeans were performed as a post-hoc test, using the emmeans’ package (Lenth, 2021) and the 5th and 95th quartiles, to test the effect of the interaction of body temperature and substrate on the reflectance of the individual.

To assess the differences in individual reflectance between night and day, we conducted a 1-factor repeated measures ANOVA. Reflectance (*n* = 208) was the dependent variable and session (*n* = 7) was the factor. The ANOVA included the reflectance measurements determined during four nights and three days (Supporting Information Figure S1). To account for the violation of the sphericity assumption, the Greenhouse-Geisser correction was applied to the 1-factor repeated measures ANOVA. Post-hoc Student’s t-tests were then used to identify significant differences in reflectance between each session.

### Software for individual recognition

#### Wild-ID®

Wild-ID (Bolger *et al*., 2012) uses the SIFT algorithm (Lowe, 2004) to identify distinctive features in each greyscale transformed image. It then evaluates pattern similarity for each pair of images by comparing the arrangement of SIFT features. A similarity score is then computed based on how well the features of the two images fit. Wild-ID provides the observer with 20 candidates, the best being the one with the highest score (a score of 1 indicating a perfect match). It is recommended to crop the images on the body area of interest with another software in order to eliminate spurious features in the background. Thus, for the use of Wild-ID, dorsal photographs of geckos were cropped using XnView 2.49.3 software (XnSoft Corp.).

#### Hotspotter®

Hotspotter (Crall *et al*., 2013) uses two algorithms. First, a one-versus-one algorithm, comparable to SIFT, extracts SIFT features based on RootSIFT (Arandjelović & Zisserman, 2012) and the Hessian-Hessian operator (Perd’och *et al*., 2009). Second, a one-versus-many algorithm employs LNBNN methods to identify images with similar groupings of features. While the one-versus-one matching algorithm compares the query image against each database image sequentially, the one-versus-many matching algorithm compares each descriptor from the query image against all descriptors from the image database. Based on the results of both algorithms, Hotspotter assigns a similarity score to each match. By default, it proposes five candidates (which can be adjusted by the user), the best candidate having the highest score. Unlike Wild-ID, Hotspotter includes a tool for cropping the images on the region of interest (ROI), then called “chips”.

For both Wild-ID and Hospotter, the user must visually compare the suggested candidates and define their status (i.e. whether or not the candidate corresponds to the investigated individual).

### Wild-ID and Hotspotter evaluation procedure

To evaluate the effect of chromatic variation on the performance of the two software packages, the analyses were carried out using dorsal side photographs with maximum and minimum reflectance differences. Two photographic databases were thus created for each of the 30 individuals.

The first database was made to assess the software’s ability to recognise the individual when the reflectance difference was at a minimum. Such database was composed of 249 photographs of individual European leaf-toed geckos collected prior to this study in various localities (Riou, Frioul and Lerins archipelagos) to which 29 photographs corresponding to the remaining 29 individuals from this study were added. The two pictures for which the reflectance difference was minimum for the investigated gecko were finally added to the database, for a total of 280 photographs.

The second database was made to assess the software’s ability to recognise the individual when the reflectance difference is at a maximum. Such database was created in the same process as above except the two provided pictures for the investigated individual displayed a maximum reflectance difference. For all 30 tested individuals, this process was repeated and the rank and similarity score were obtained.

The error rates were quantified with False Rejection Rate (FRR) and calculated for the top ranking candidate. The FRR corresponds to the probability that a recapture event is falsely identified as a new capture (i.e. the software fails to match two photos of the same individual). FRR is calculated as the ratio between the number of false rejections and the total number of identification attempts, and thus ranges from 0 (i.e. 100% success in matching two different images) to 1 (0% success).

Because data failed to approximate normal distribution, a non-parametric Wilcoxon-Mann-Whitney test was used to evaluate the differences between the similarity scores obtained by the same software in conditions of maximum and minimum reflectance differences. The comparison of the similarity scores between Wild-ID and Hotspotter could not be performed because each software has a specific method of calculating the scores.

For all analyses, when necessary, normality of residuals and homoscedasticity were checked. All analyses were performed using R software (v.4.1.0) (R Core Team, 2021). All plots were generated with the ggplot2 package (Wickham, 2016).

## Results

### Reflectance spectrum and factors influencing chromatic variation

The dorsal reflectance of the 30 European leaf-toed geckos varied between a minimum of 0.088 and maximum of 0.310 (median = 0.190). The mean reflectance was 0.189 ± 0.047 SD.

Based on AIC scores of the GLMM, a model that included substrate type, body temperature and the interaction between substrate type and body temperature was the top ranked model (Table 1). The selected model showed that the dorsal side of the geckos have a significantly lower reflectance in the vegetated micro-habitat (VMH) than in the rocky micro-habitat (RMH) for both low (T = 14.0°C; EMMEans pairwise comparison: *P* < 0.05) and high body temperatures (T = 25.6°C; EMMEans pairwise comparison: *P* < 0.001). We also observed that this difference increases with higher body temperature, with reflectance decreasing more strongly on vegetated substrates for the same given body temperature (Fig. 1).

**Table 1.** Summary of the best-fitting model (dAICc < 2) and null model for the effect of the body temperature and the substrate used on European leaf-toed gecko reflectance. Marginal (Rm^2^) and conditional (Rc^2^) R^2^ values are indicating the goodness-of-fit of the models.

| Response variable | df | AICc | dAICc | Akaike weight (wi) | Explanatory variable | $R_m^2$ | $R_c^2$ |
| --- | --- | --- | --- | --- | --- | --- | --- |
| Reflectance | 7 | -662.70 | 0.00 | 1 | BodyT+Substrate+<br>BodyT:Substrate | 0.746 | NA |
| Reflectance | 4 | -500.20 | 162.44 | 0 | 1 | 0.000 | 0.258 |

**Figure 1.**
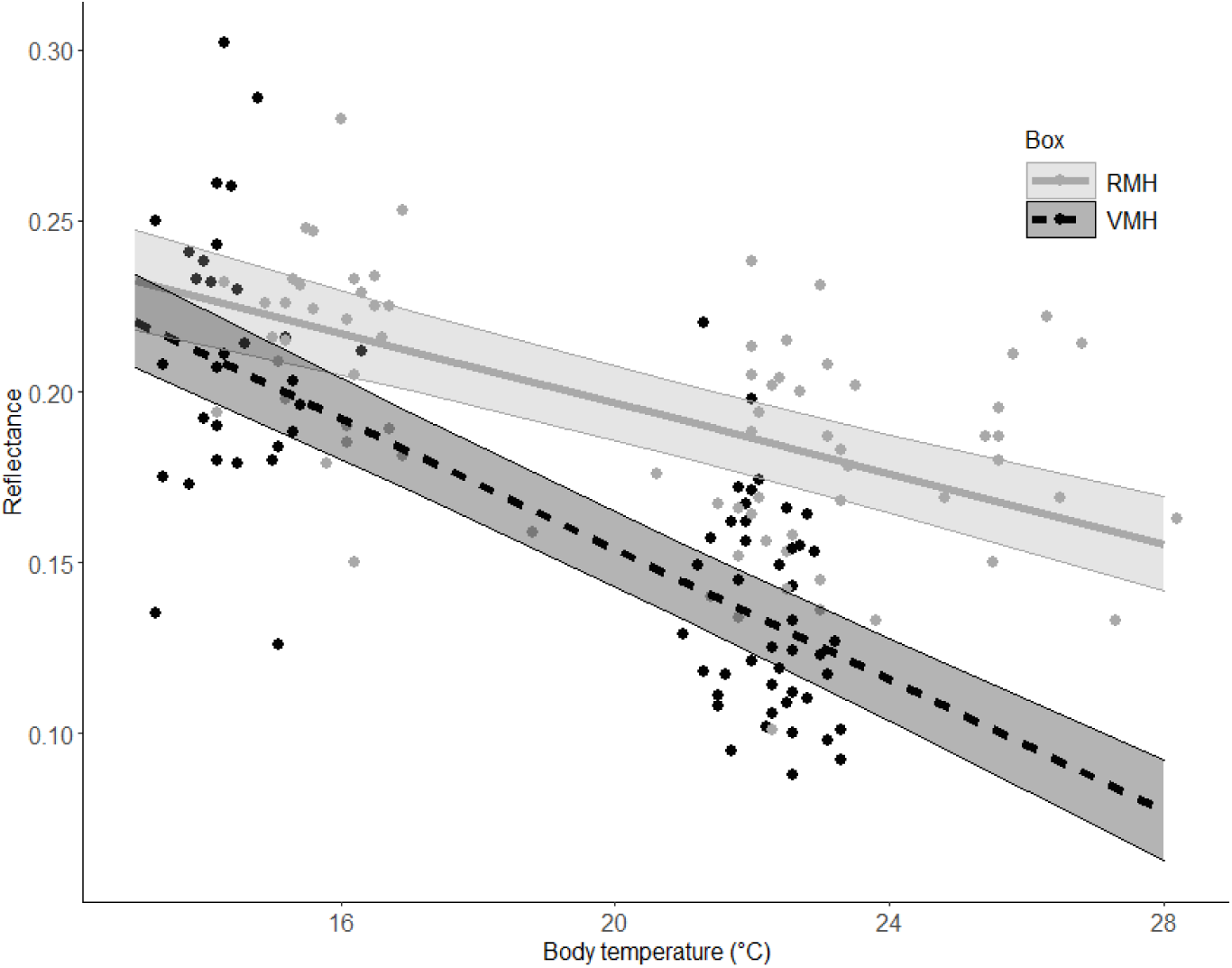
Body temperature effect on the dorsal reflectance of the European leaf-toed geckos for the two substrate types used. The solid line represents data collected on light-coloured rocky substrate (RMH) while the dashed line represents dark-coloured vegetated substrate (VMH) data. Predicted mean value and 95% confidence intervals are displayed for each substrate type and based on the best fitting model selected.

The repeated measures 1-factor ANOVA showed that the dorsal side of individuals exhibited a significantly different reflectance as a function of session (*F*_2.58,69.72_ = 33.68 ; *P* < 0.001). Reflectance is significantly lower in daytime sessions than in nighttime sessions (Post-hoc Student’s t test, *P* < 0.001) (Fig. 2). In contrast, reflectance was not significantly different between daytime sessions or between nighttime sessions (Post-hoc Student’s t test, *P* > 0.05).

**Figure 2.**
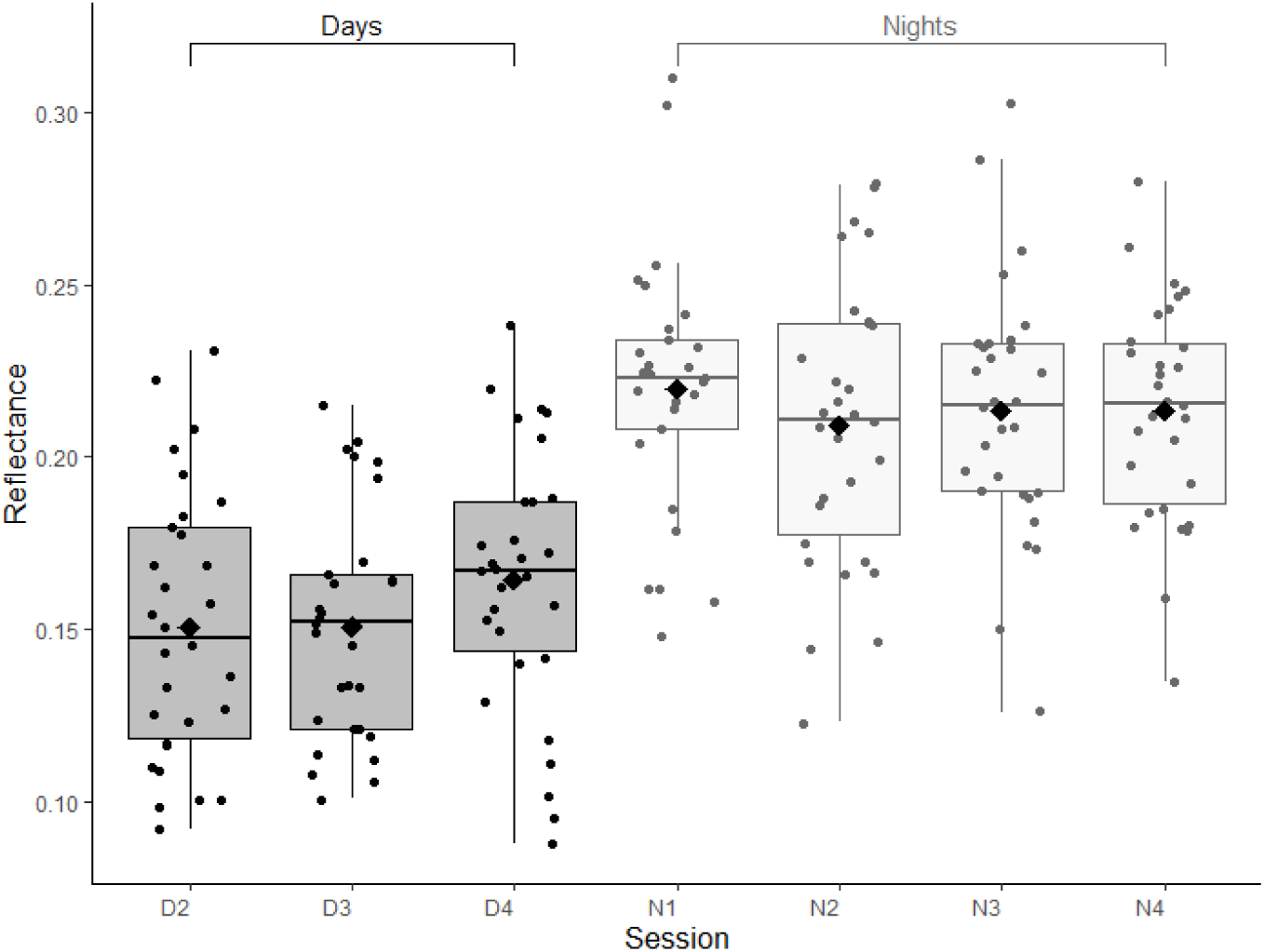
Boxplots displaying the effect of the different sessions on the dorsal reflectance of the European leaf-toed geckos. Sessions N1, N2, N3 and N4 were conducted at night ; sessions D2, D3 and D4 were conducted during the day. The first and third quartiles are represented by the bottom and the top of the boxes respectively. The middle line corresponds to the median and the black diamonds the reflectance averages. The minimum and maximum values are represented with the vertical axes. Beyond that, the points represent outliers.

### Effects of chromatic variation and software on FRRs

For Wild-ID and Hotspotter, the top ranked candidate was always the true matching image for both minimum and maximum reflectance differences (FRR = 0). Thus both software packages correctly recognised the image pairs for each individual.

For Wild-ID, the scores obtained for maximum reflectance differences (mean: 0.143 ± 0.090 SD) were significantly lower than minimum reflectance differences (mean: 0.333 ± 0.182 SD) (Wilcoxon-Mann-Whitney, W = 165, *P* < 0.001) (Fig. 3a). Conversely, for Hotspotter, the similarity scores obtained for maximum reflectance differences (mean: 364.933 ± 170.065 SD) were not significantly different from the scores obtained for minimum reflectance differences (mean: 506.867 ± 297.575 SD) (Wilcoxon-Mann-Whitney, W = 321.5, *P* = 0.057) (Fig. 3b).

**Figure 3.**
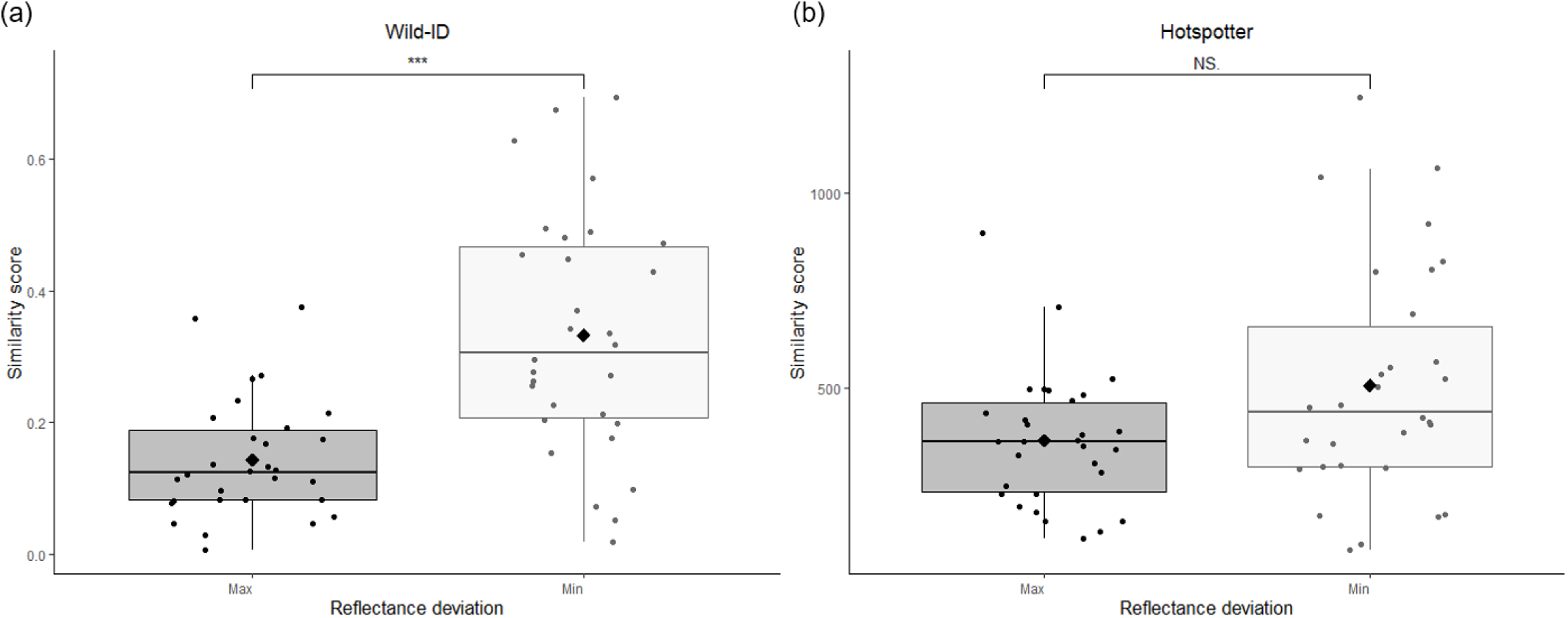
Similarity scores obtained for (a) Wild-ID and (b) Hotspotter in maximum (Max) and minimum (Min) reflectance difference conditions. NS = Not Significant, *** = Significant (*P* < 0.001). Black diamonds represent the means of the similarity scores. The higher the score, the better the correspondence between two photographs.

## Discussion

The extent of individual colour variation for the European leaf-toed gecko was characterised based on reflectance values extracted from animals’s dorsal photographs. Parameters involved in the colouration changes of the species were then identified in a semi-controlled environment. Finally, the applicability and reliability of two individual recognition software programs were evaluated using European leaf-toed geckos dorsal side photographs for both maximum and minimum reflectance differences for each individual monitored. We found that (1) the colouration of the European leaf-toed gecko was influenced by the combined effect of the type of substrate of their environment and their body temperature. Individuals were also significantly darker during the day than at night, regardless of the environment type in which they were found. (2) Based on the top ranking candidate, Wild-ID and Hotspotter correctly identified 100% of matching image pairs. (3) Maximum reflectance differences did not significantly decrease similarity scores for Hotspotter, unlike Wild-ID.

### Physiological colour change in the European leaf-toed gecko

Our results show that the European leaf-toed gecko is able to adjust its dorsal reflectance as a response to different environmental conditions. On a darker substrate, the reflectance values obtained were significantly lower (i.e. darker individuals) than on a lighter substrate whether by day or by night. The effect of background on colouration has been previously demonstrated in two sympatric gecko species (*Hemidactylus turcicus, Tarentola mauritanica*) (Zaidan & Wiebusch, 2007; Vroonen *et al*., 2012). The phenomenon of homochromia, which makes individuals less visible in their habitat, is known to be an adaptation reducing predation risk in saurians (Stuart-Fox, Whiting & Moussalli, 2006, Stuart-Fox & Moussali, 2008; Ito, Ikeuchi & Mori, 2013). It is therefore highly probable that the European leaf-toed gecko exhibits a similar evolutionary mechanism.

Pigmentary cells are also likely to be influenced by the body temperature of individuals, the reflectance values being lower as body temperature increases. As the European leaf-toed gecko spends most of its daytime hidden in rock crevices, a darker colouration might allow a better assimilation of solar radiation. It was shown for many ectotherms that a darker colouration might allow quicker reaching of an optimal body temperature compared to lighter colouration (Walton & Bennett, 1993; Silbiger & Munguia, 2008; Langkilde & Boronow, 2012; Kronstadt *et al*., 2013; Smith *et al*., 2016). However, a significant difference in reflectance between night and day (i.e. individuals were darker during the day than at night) also indicates a potential effect of light in the European leaf-toed gecko skin colouration. Indeed, some authors showed that under controlled conditions, light could trigger melanin dispersion in the geckos *Hemidactylus turcicus* (Zaidan & Wiebusch, 2007) and *Tarentola mauritanica* (Vroonen *et al*., 2012). However, the semi-controlled conditions of our experiment did not allow us to dissociate the effect of body temperature and the effect of light, both of which are influenced by natural day/night cycles.

All these elements suggest that the important variation in colour patterns is induced by complex interactions between the substrate type, the body temperature which is highly correlated to the substrate temperature, as well as the probable influence of light.

### Consequences of chromatic variation on algorithmic performance

With a 100% success rate (FRR = 0) in the top ranking candidate match, Wild-ID and Hotspotter proved to be extremely reliable in recognising European leaf-toed geckos. Despite the large reflectance differences, both software programs were highly successful in identifying SIFT features and matching them (Fig. 4a and 4b). By performing a series of image transformations including greyscale conversion, contrast and brightness adjustment (Lowe, 2004), SIFT algorithm allows the dorsal patterns (i.e. pale dorsal line and transverse bands) to reappear. Thus, the SIFT algorithm seems to identify the contours of the patterns by selecting the darkest scales, contrasting with the light dorsal patterns. Different studies showed that SIFT algorithm relies on the most contrasting patterns of the photograph (i.e. displaying a sharp or graded break in multiple directions) that are easily localisable (e.g. giraffe spot contours, zebra stripe contours) (Bolger *et al*., 2012; Crall *et al*., 2013). Since scale insertions in the European leaf-toed gecko display contrasting patterns between coloured (lighter insertions) and uncoloured (darker insertions) scales, individuals could be more easily recognised by the algorithm, potentially explaining this very high recognition rate of both software on the species.

**Figure 4.**
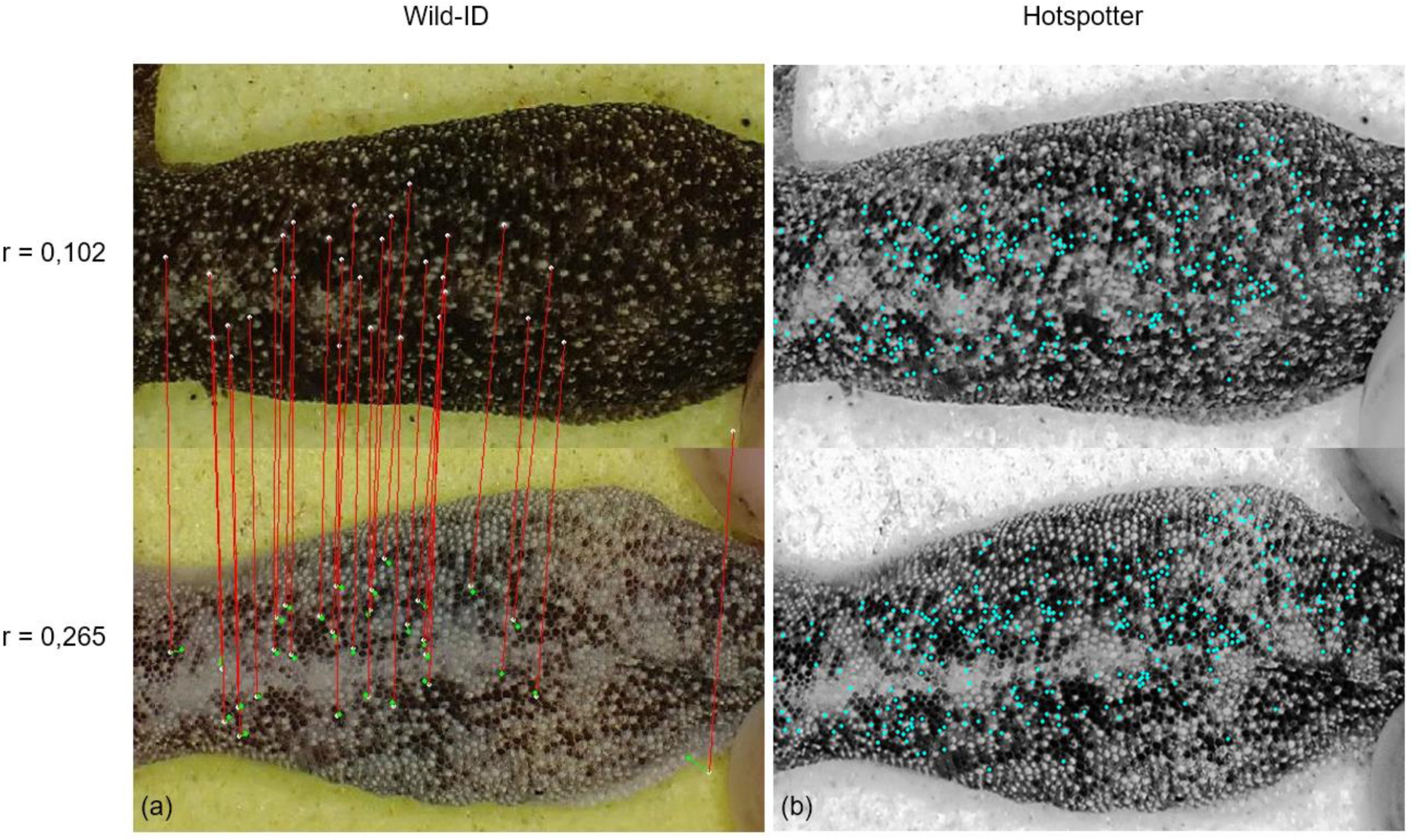
The SIFT features extracted by (a) Wild-ID (white points) and (b) Hotspotter (blue points) that are similar for the image pair of the individual 13 in maximum reflectance difference condition. The reflectance (r) is specified for the investigated individual (top) and the suggested candidate by the software (bottom). The red lines connect the SIFT features identified by Wild-ID and the green points correspond to the location where the SIFT features of the suggested candidate should be after affine transformation of the investigated individual. Note that Wild-ID does not provide a visual of the greyscale images.

Therefore, we assume that saurian species capable of rapid colour change and characterised by horny scales or tubercles with an individual and stable arrangement over time (Steinicke *et al*., 2000; Perera & Perez-Mellado, 2004) might be good candidates for computer-assisted photo-identification (Fig. 5).

**Figure 5.**
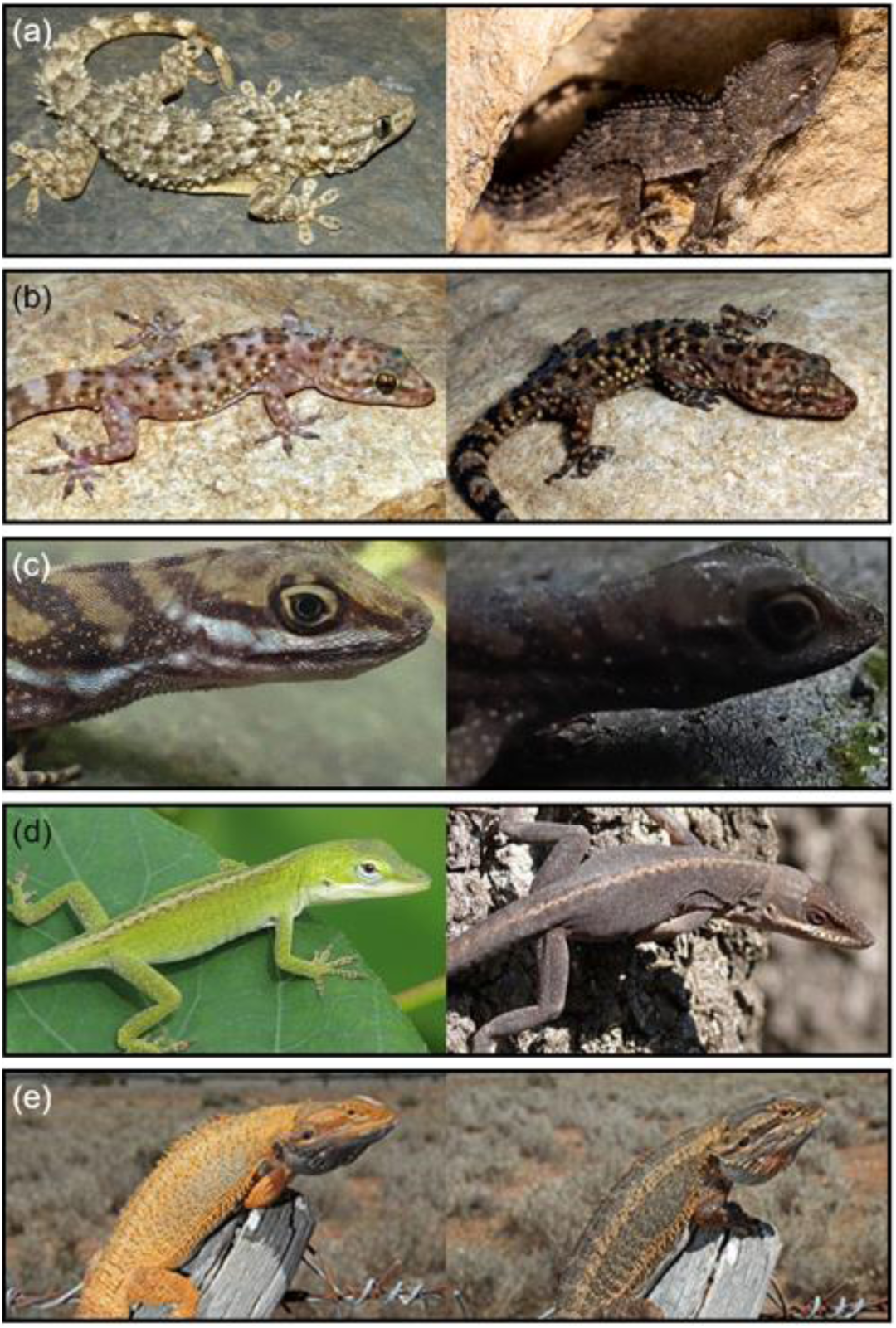
Example of saurians species able to colour change. (a) *Tarentola mauritanica*. Photo credit : Daniel Jablonski (left) and Jacques Jean (right). (b) *Hemidactylus turcicus*. Photo credit : Gary Nafis. (c) *Anolis aquaticus*. Photo credit : Lindsey Swierk. (d) *Anolis carolinensis* Photo credit : Vicki DeLoach / Woodstock, Georgia USA (left) and Will Cook (right). (e) *Pogona vitticeps*. Photo credit : Adam Elliot.

Similarity score analysis further revealed small variations in the performance of the two software packages. Wild-ID significantly displayed lower similarity scores for maximum reflectance differences compared to minimum reflectance differences. In contrast, Hotspotter’s similarity scores did not differ significantly between the two reflectance conditions. This finding indicates that the Hotspotter software is less sensitive to chromatic variation than Wild-ID, and therefore may be less likely to generate false rejection errors as sample sizes of unknown individuals increase. The combination of the two algorithms, one-versus-one and one-versus-many, probably provides Hotspotter with a highly reliable identification procedure. Such performance differences between the two software (Wild-ID vs Hotspotter) were also attested by several authors which have already concluded that Hotspotter performed better than Wild-ID (Crall *et al*., 2013; Chehrsimin *et al*., 2018; Nipko *et al*., 2020).

### Management implications

When monitoring species with computer-assisted photo-identification, the standardisation of photos is imperative to obtain sharp and high quality images (Kelly, 2001; Bendik *et al*., 2013; Morrison *et al*., 2016; Sacchi *et al*., 2016) which will be more likely to match (Nipko *et al*., 2020). For this purpose, the use of a portable “photographic studio” comparable to the one suggested here for monitoring a small gecko (Supporting Information Figure S2) is essential to ensure safe handling and stability of photographic parameters (angle of view, distance between the animal and the camera, non-reflected light, etc.). With such a device, two people are enough to hold and photograph the individuals in the field. Within the framework of a capture-recapture study of the European leaf-toed gecko, we advise to carry out the photographs at night to conform with the activity time of this species.

Hotspotter offers more options and tools to ease the pre-processing of images and their analysis (Nipko *et al*., 2020). The creation of “chips” allows for quick extraction of the region of interest from the photographs being compared without relying on any external software. The orientation of the animal can also be defined, allowing oblique postures. Users can visualise the SIFT features on which Hotspotter relies to compare images. Such software can integrate individuals over the course of capture-recapture sessions without the need to reprocess old photographs, unlike Wild-ID. The user can label identified individuals for referencing. The automated greyscale conversion of the chips makes the visual recognition of European leaf-toed geckos possible and allows the user to make the final decision (i.e. if the suggested candidate by the software corresponds or not to the investigated individual). In light of all these advantages, we highly recommend the use of Hotspotter for the demographic monitoring of the European leaf-toed geckos. A better understanding of the population dynamics will allow the implementation of relevant conservation actions for this species.

## Conclusions

After investigating parameters causing the physiological colour change of the European leaf-toed gecko, we showed for the first time that high chromatic variations had no impact on the reliability of the two most widely used free software programs for individual wildlife recognition. Hotspotter proved to be less sensitive to colour changes and its interface is more suitable for species monitoring. These results should encourage biologists to use this non-invasive technology for individual monitoring of the European leaf-toed gecko and other saurian species capable of rapid colour change in their natural environment. Further studies on smooth-skinned taxa that change colour (e.g. amphibians or cephalopods) would allow a better assessment of the role of skin structure in computer-assisted individual recognition. At the same time, the results will help to appreciate the extent of the possible use of these tools for non-invasive wildlife monitoring.

## Acknowledgements

We warmly thank Amanda Xérès (CEN PACA), Félix Thirion (CEN PACA), Nathan Huvier, Corentin Lambert-Grimpard (Muséum d’Histoire Naturelle de Nice), Alexandre Viguier (Office Français de la Biodiversité) for their assistance in the field and Olivier Gerriet (Muséum d’Histoire Naturelle de Nice) who designed the terrariums and helped us on the field. This study was made possible thanks to the logistical support of the Office National des Forêts (Eric Tassone) and the Parc National des Calanques (Lidwine Le Mire Pecheux, Lorraine Anselme, Alain Mante and Timothée Cuchet). We also thank Margaux Treguy for her precious help on the use of the GLMMs and GDM, Thomas Morrison (University of Glasgow), Jonathan P. Crall (Kitware Inc.), Agata Staniewicz (University of Bristol), Maximilian Matthé (University of Dresden), Hugo Cayuela (Université de Lyon) and Giacomo Rosa (University of Genoa) for providing useful comments which helped to significantly improve the quality of the manuscript. The English translation was assisted by Félix Thirion, whom we sincerely thank. This work was funded by the Provence-Alpes-Côte d’Azur region and supported by the Parc National des Calanques. This study has been authorised by the French government by prefectural decree for the departments of Alpes-Maritimes (permit number : 2020_341) and Bouches-du-Rhône (permit number : 2020_250 and 2021_113).

## Conflict of interest

No conflict of interest to declare.

## Author’s contributions

JR and CM conceptualised and designed the study. JR supervised the field data collection and the overall study. CM and TD carried out the data analysis and participated in the field data collection. JR, CM and TD wrote the first manuscript draft. All authors revised the manuscript.

## Notes

### Competing Interest Statement

The authors have declared no competing interest.

## References

Arandjelovic, R. & Zisserman, A. (2012). Three things everyone should know to improve object retrieval. In 2012 IEEE Conference on Computer Vision and Pattern Recognition: 2911-2918. Providence.

Bagnara, J. T. & Hadley, M. E. (1973). Chromatophores and Color Changes: The Comparative Physiology of Animal Pigmentation. Englewood Cliffs: Prentice-Hall.

Bardier, C., Székely, D., Augusto-Alves, G., Matínez-Latorraca, N., Schmidt, B. R. & Cruickshank, S. S. (2020). Performance of visual vs. software-assisted photo-identification in mark-recapture studies: a case study examining different life stages of the Pacific Horned Frog (Ceratophrys stolzmanni). Amphib.-Reptil. 42, 17–28.

Barord, G. J., Dooley, F., Dunstan, A., Ilano, A., Keister, K. N., Neumeister, H., Preuss, T., Schoepfer, S. & Ward, P. D. (2014). Comparative population assessments of Nautilus sp. in the Philippines, Australia, Fiji, and American Samoa using baited remote underwater video systems. PLoS One 9, e100799.

Barton, K. (2020). MuMIn: Multi-Model Inference. R package version 1.43.17. Available online: https://CRAN.R-project.org/package=MuMIn. Accessed on 07 July 2021.

Baskale, E. & Kaya, U. (2012). Decline of the Levantine Frog, Pelophylax bedriagae Camerano, 1882, in the western Aegean Region of Turkey changes in population size and implications for conservation: (Amphibia: Ranidae). Zool. Middle East 57, 69–76.

Bendik, N. F., Morrison, T. A., Gluesenkamp, A. G., Sanders, M. S. & O’Donnell, L. J. (2013). Computer-assisted photo identification outperforms visible implant elastomers in an endangered salamander, Eurycea tonkawae. PloS One 8, e59424.

Bengsen, A., Butler, J. & Masters, P. (2011). Estimating and indexing feral cat population abundances using camera traps. Wildl. Res. 38, 732–739.

Bolger, D. T., Morrison, T. A., Vance, B., Lee, D. & Farid, H. (2012). A computer-assisted system for photographic mark–recapture analysis. Methods Ecol. Evol. 3, 813–822.

Boyer, J. F. & Swierk, L. (2017). Rapid body color brightening is associated with exposure to a stressor in an Anolis lizard. Can. J. Zool. 95, 213–219.

Burnham, K. P. and Anderson, D. R (2002). Model selection and multimodel inference: a practical information-theoretic approach. 2nd edn. New York: Springer.

Chehrsimin, T., Eerola, T., Koivuniemi, M., Auttila, M., Levänen, R., Niemi, M., Kunnasranta, M. & Kälviäinen, H. (2018). Automatic individual identification of Saimaa ringed seals. IET Comput. Vis. 12, 146–152.

Corti, C., Cheylan, M., Geniez P., Sindaco, R. & Romano, A. (2009). Euleptes europaea (Gené, 1839). The IUCN Red List of Threatened Species 2009: e.T61446A12486542. Version 2021-3. https://dx.doi.org/10.2305/IUCN.UK.2009.RLTS.T61446A12486542.en. Accessed on 13 January 2022.

Costa, A., Oneto, F., & Salvidio, S. (2019). Time-for-space substitution in N-mixture modeling and population monitoring. J. Wildl. Manage. 83, 737–741.

Crall, J. P., Stewart, C. V., Berger-Wolf, T. Y., Rubenstein, D. I. & Sundaresan, S. R. (2013). Hotspotter – Patterned species instance recognition. In 2013 IEEE Workshop on Applications of Computer Vision (WACV): 230–237. Florida.

Cross, M. D., Lipps Jr, G. J., Sapak, J. M., Tobin, E. J. & Root, K. V. (2014). Pattern-recognition software as a supplemental method of identifying individual eastern box turtles (Terrapene c. carolina). Herpetol. Rev. 45, 584–586.

Cruickshank, S. S. & Schmidt, B. R. (2017). Error rates and variation between observers are reduced with the use of photographic matching software for capture-recapture studies. Amphib.Reptil. 38, 315–325.

Delaugerre, M. (1981a). Sur l’histoire naturelle de Phyllodactylus europaeus Gené, 1838 (Gekkonidae Sauria Reptiles). Port-Cros : étude d’une population naturelle. Trav. Sci. Parc Nation. Port-Cros 6, 147–175.

Delaugerre, M. (1981b). Un cas d’albinisme chez Phyllodactylus europaeus Gene, 1838. Premier cas signalé dans la famille des Gekkonidae (Sauria-Reptiles). Bull. Mens. Soc. Linn. Lyon 50, 213–216.

Delaugerre, M. (1992). Le phyllodactyle d’Europe Phyllodactylus europaeus Gené, 1839. In Atlas de Répartition des Batraciens et Reptiles de Corse: 60–63.

Delaugerre, M. & Cheylan, M. (Eds). Ajaccio, Montpellier: Parc Naturel Régional & Ecole Pratique des Hautes Etudes.

Delaugerre, M. (1997). Phyllodactylus europaeus. In Atlas of Amphibians and Reptiles in Europe: 212–213.

Gasc, J.-P., Cabela, A., Crnobrnja-isailovic, J., Dolmen, D., Grossenbacher, K., Haffner, P., Lescure, J., Martens, H., Martìnez Rica, J.-P., Maurin, H., Oliveira, M.-E., Sofianidou, T., Veith, M. & Zuiderwijk, A. (Eds). Paris: Societas Europea Herpetologica et Muséum national d’Histoire naturelle, IEGB/SPN.

Delaugerre, M., Ouni, R. & Nouira, S. (2011). Is the European Leaf-toed gecko Euleptes europaea also an African ? Its occurrence on the Western Mediterranean landbrige islets and its extinction rate. Herpetol. Notes 4, 127–137.

Dunbar, S. G., Anger, E. C., Parham, J. R., Kingen, C., Wright, M. K., Hayes, C. T., Safi, S., Holmberg, J., Salinas, L. & Baumbach, D. S. (2021). HotSpotter: Using a computer-driven photo-id application to identify sea turtles. J. Exp. Mar. Biol. Ecol. 535, 151490.

Endler, J. A. (1990). On the measurement and classification of colour in studies of animal colour patterns. Biol. J. Linn. Soc. 41, 315–352.

Fan, M., Stuart-Fox, D. & Cadena, V. (2014). Cyclic colour change in the bearded dragon Pogona vitticeps under different photoperiods. PloS One 9, e111504.

Hadley, M. E & Goldman, J. (1969). Physiological Color Change in Reptiles. Am. Zool. 9, 489–504.

Hamilton, P. S., Gaalema, D. E., Laage, S. L. & Sullivan, B. K. (2005). A photographic method for quantifying color characteristics and color patch dimensions in lizards. Herpetol. Rev. 36, 402.

Hamilton, P., Gaalema, D. E. & Sullivan, B. (2008). Short-term changes in dorsal reflectance for background matching in Ornate Tree Lizards (Urosaurus ornatus). Amphib.-Reptil. 29, 473–477.

Ikeda, Y. (2021). Color Change in Cephalopods. In Pigments, Pigment Cells and Pigment Patterns: 425–449.

Hashimoto, H., Goda, M., Futahashi, R., Kelsh, R. & Akiyama, T. (Eds). Singapore: Springer.

Ito, R., Ikeuchi, I. & Mori, A. (2013). A day gecko darkens its body color in response to avian alarm calls. Curr. Herpetol. 32, 26–33.

Jackson, R. M., Roe, J. D., Wangchuk, R. & Hunter, D. O. (2006). Estimating snow leopard population abundance using photography and capture-recapture techniques. Wildl. Soc. Bull. 34, 772–781.

Jain, A. K. (2007). Biometric recognition. Nature, 449, 38–40.

Johnson, J.W. (2000). A Heuristic Method for Estimating the Relative Weight of Predictor Variables in Multiple Regression. Multivar. Behav. Res. 35, 1–19.

Kang, C., Kim, Y. E. & Jang, Y. (2016). Colour and pattern change against visually heterogeneous backgrounds in the tree frog Hyla japonica. Sci. Rep. 6, 1–12.

Kelly, M. J. (2001). Computer-aided photograph matching in studies using individual identification: an example from Serengeti cheetahs. J. Mammal. 82, 440–449.

Kindermann, C. & Hero, J. M. (2016). Pigment cell distribution in a rapid colour changing amphibian (Litoria wilcoxii). Zoomorphology, 135, 197–203.

Kronstadt, S. M., Darnell, M. Z. & Munguia, P. (2013). Background and temperature effects on Uca panacea color change. Mar. Biol. 160, 1373–1381.

Langkilde, T. & Boronow, K. E. (2012). Hot boys are blue: temperature-dependent color change in male eastern fence lizards. J. Herpetol. 46, 461–465.

Lea, J. M., Walker, S. L., Kerley, G. I., Jackson, J., Matevich, S. C. & Shultz, S. (2018). Non-invasive physiological markers demonstrate link between habitat quality, adult sex ratio and poor population growth rate in a vulnerable species, the Cape mountain zebra. Funct. Ecol. 32, 300–312.

Lenth, R.V. (2021). Emmeans: estimated marginal means, aka least-squares means. R package version 1.5.4. Available online: https://CRAN.R-project.org/package=emmeans. Accessed on 07 July 2021.

Lewis, A. C., Rankin, K. J., Pask, A. J. & Stuart-Fox, D. (2017). Stress-induced changes in color expression mediated by iridophores in a polymorphic lizard. Ecol. Evol. 7, 8262–8272.

Ligon, R. A. & McGraw, K. J. (2013). Chameleons communicate with complex colour changes during contests: different body regions convey different information. Biol. Lett. 9, 20130892.

Ligon, R. A. (2014). Defeated chameleons darken dynamically during dyadic disputes to decrease danger from dominants. Behav. Ecol. Sociobiol. 68, 1007–1017.

Lowe, D. G. (2004). Distinctive image features from scale-invariant keypoints. Int. J. Comput. Vis. 60, 91–110.

Lüdecke, D., Ben-Shachar, M., Patil, I., Waggoner, P. & Makowski, D. (2021). Performance: An R Package for Assessment, Comparison and Testing of Statistical Models. J. Open Source Softw. 6, 3139.

Magnusson, A., Skaug, H., Nielsen, A., Berg, C., Kristensen, K., Maechler, M., van Bentham, K., Bolker, B., Brooks, M. & Brooks, M. M. (2017). Package ‘glmmTMB’. R Package Version 1.1.1. Available online: https://CRAN.R-project.org/package=glmmTMB. Accessed on 07 July 2021.

Marchand, M. A., Roy, C., Renet, J., Delauge, J., Meyer, D. & Hayot, C. (2017). Liste rouge régionale des amphibiens et reptiles de Provence-Alpes-Côte d’Azur. 14p.

Matthé, M., Sannolo, M., Winiarski, K., Spitzen-van der Sluijs, A., Goedbloed, D., Steinfartz, S. & Stachow, U. (2017). Comparison of photo-matching algorithms commonly used for photographic capture–recapture studies. Ecol. Evol. 7, 5861–5872.

McCann, S. & Lowe, D. G. (2012). Local Naive Bayes Nearest Neighbor for image classification. In 2012 IEEE Conference on Computer Vision and Pattern Recognition: 3650–3656. Providence.

Morrison, T. A., Yoshizaki, J., Nichols, J. D. & Bolger, D. T. (2011). Estimating survival in photographic capture–recapture studies: overcoming misidentification error. Methods Ecol. Evol. 2, 454–463.

Morrison, T. A. & Bolger, D. T. (2014). Connectivity and bottlenecks in a migratory wildebeest Connochaetes taurinus population. Oryx 48, 613–621.

Morrison, T. A., Keinath, D., Estes-Zumpf, W., Crall, J. P. & Stewart, C. V. (2016). Individual identification of the endangered Wyoming toad Anaxyrus baxteri and implications for monitoring species recovery. J. Herpetol. 50, 44–49.

Nakagawa, S., Johnson, P. C. D. & Schielzeth, H. (2017). The coefficient of determination R2 and intra-class correlation coefficient from generalized linear mixed-effects models revisited and expanded. J. R. Soc. Interface 14, 20170213.

Nipko, R. B., Holcombe, B. E. & Kelly, M. J. (2020). Identifying Individual Jaguars and Ocelots via Pattern-Recognition Software: Comparing HotSpotter and Wild-ID. Wildl. Soc. Bull. 44, 424–433.

Pace, R. M. III, Corkeron, P. J. & Kraus, S. D. (2017). State–space mark–recapture estimates reveal a recent decline in abundance of North Atlantic right whales. Ecol. Evol. 7, 8730–8741.

Perd’och, M., Chum, O. & Matas, J. (2009). Efficient representation of local geometry for large scale object retrieval. In 2009 IEEE Conference on Computer Vision and Pattern Recognition: 9–16. Miami.

Perera, A. & Perez-Mellado, V. (2004). Photographic identification as a non invasive marking technique for Lacertid lizard. Herpetol. Rev. 35, 349–350.

Quinby, B. M., Creighton, J. C. & Flaherty, E. A. (2021). Estimating Population Abundance of Burying Beetles Using Photo-Identification and Mark-Recapture Methods. Environ. Entomol. 50, 238–246.

R Core Team (2021). R: A language and environment for statistical computing. Vienna, Austria: R Foundation for Statistical Computing.

Renet, J., Gerriet, O., Kulesza, V. & Delaugerre, M. (2013). Le Phyllodactyle d’Europe Euleptes europaea (Gené, 1839) (Reptilia, Squamata, Sphaerodactylidae)—Les populations continentales françaises ont-elles un avenir ?. Bull. Soc. Herp. Fr. 145, 189–198.

Renet, J., Leprêtre, L., Champagnon, J. & Lambret, P. (2019). Monitoring amphibian species with complex chromatophore patterns: a non-invasive approach with an evaluation of software effectiveness and reliability. Herpetol. J. 29, 13–22

Sacchi, R., Scali, S., Mangiacotti, M., Sannolo, M. & Zuffi, M. A. (2016). Digital identification and analysis. In Reptile Ecology and Conservation. A Handbook of Techniques: 59–72. C. K. Dodd. (Ed). New-York: Oxford University Press.

Schneider, C. A., Rasband, W. S. & Eliceiri, K. W. (2012). NIH Image to ImageJ: 25 years of image analysis. Nat. Methods 9, 671–675.

Schofield, G., Klaassen, M., Papafitsoros, K., Lilley, M. K., Katselidis, K. A. & Hays, G. C. (2020). Long-term photo-id and satellite tracking reveal sex-biased survival linked to movements in an endangered species. Ecology 101, e03027.

Silbiger, N. & Munguia, P. (2008). Carapace color change in Uca pugilator as a response to temperature. J. Exp. Mar. Biol. Ecol. 355, 41–46.

Sköld, H. N, Aspengren, S. & Wallin, M. (2013). Rapid color change in fish and amphibian—function, regulation, and emerging applications. Pigment Cell Melanoma Res. 26, 29–38.

Sköld, H. N., Aspengren, S., Cheney, K. L. & Wallin, M. (2016). Fish chromatophores—from molecular motors to animal behavior. Int. Rev. Cell. Mol. Biol. 321, 171–219.

Smith, K. R., Cadena, V., Endler, J. A., Porter, W. P., Kearney, M. R. & Stuart-Fox, D. (2016). Colour change on different body regions provides thermal and signalling advantages in bearded dragon lizards. Proc. Royal Soc. B 283, 20160626.

Steinicke, H., Ulbrich, K., Henle, K. & Grosse, W. R. (2000). Eine neue Methode zur fotografischen Individualidentifikation mittelerupäischer Halsbandeidechsen (Lacertidae). Salamandra 36, 81–88.

Stuart-Fox, D., Whiting, M. J. & Moussalli, A. (2006). Camouflage and colour change: antipredator responses to bird and snake predators across multiple populations in a dwarf chameleon. Biol. J. Linn. Soc. 88, 437–446.

Stuart-Fox, D. & Moussalli, A. (2008). Selection for social signalling drives the evolution of chameleon colour change. PLoS Biol. 6, e25.

Sun, J., Wu, W., Liu, C. & Tong, J. (2017). Investigating the nanomechanical properties and reversible color change properties of the beetle Dynastes tityus. J. Mater. Sci. 52, 6150–6160.

Tabuki, K., Nishizawa, H., Abe, O., Okuyama, J. & Tanizaki, S. (2021). Utility of carapace images for long-term photographic identification of nesting green turtles. J. Exp. Mar. Biol. Ecol. 545, 151632.

Tonidandel, S. & LeBreton, J. M. (2010). Determining the Relative Importance of Predictors in Logistic Regression: An Extension of Relative Weight Analysis. Organ. Res. Methods 13, 767–781.

Vroonen, J., Vervust, B., Fulgione, D., Maselli, V. & Van Damme, R. (2012). Physiological colour change in the Moorish gecko, Tarentola mauritanica (Squamata: Gekkonidae): effects of background, light, and temperature. Biol. J. Linn. Soc. 107, 182–191.

Walton, B. M. & Bennett, A. F. (1993). Temperature-dependent color change in Kenyan chameleons. Physiol. Zool. 66, 270–287.

Wickham, H. (2016). Ggplot2: Elegant Graphics for Data Analysis. 2nd edn. New York: Springer.

Wunderlin, J. & Kropf, C. (2013). Rapid colour change in spiders. In Spider ecophysiology: 361–370.

Nentwig W. (Ed). Berlin: Springer.

Zaidan, F. & Wiebusch, P. L. (2007). Effects of temperature and illumination on background matching in Mediterranean geckos (Hemidactylus turcicus). Tex. J. Sci. 59, 127–136.

