## Supporting information for "Rapid colour changes in a tiny threatened gecko do not impede computer-assisted individual recognition"

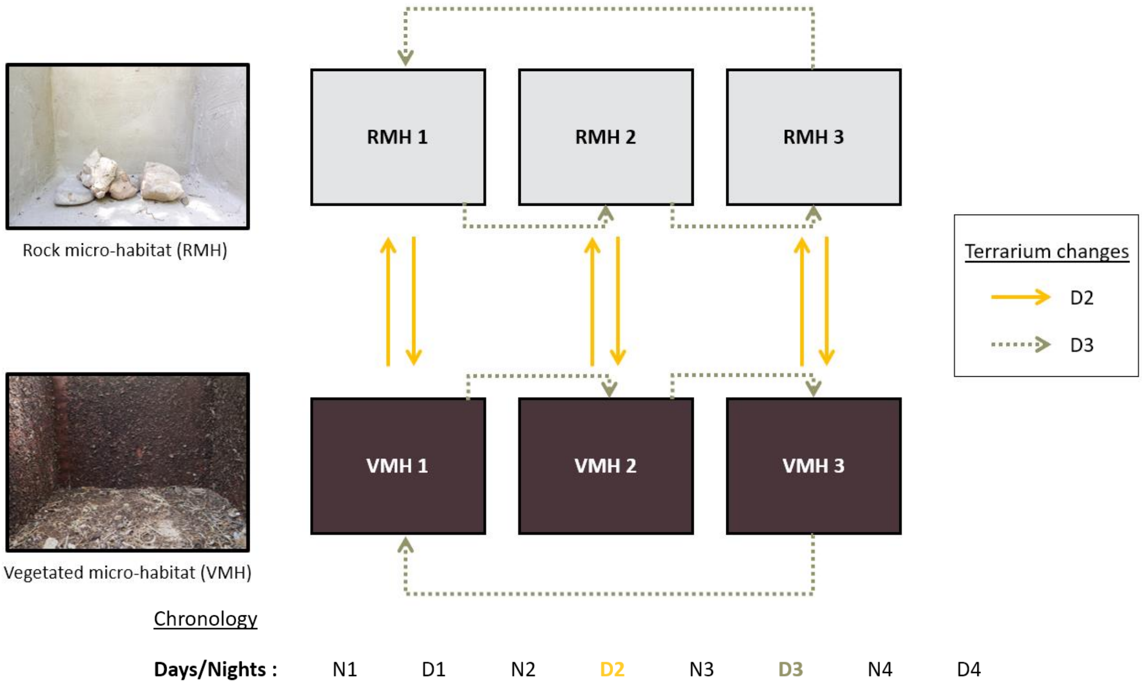


**Figure S1 :** Experimental design to study reflectance variations in the European leaf-toed gecko. This scheme presents the organisation of the terrariums in which 30 European leaf-toed geckos were maintained in captivity, distributed equally (5 individuals per terrarium), the terrarium changes of the individuals and the chronology of the night (N) and day (D) sessions. Three terrariums mimic a rocky micro-habitat (RMH) and three other terrariums mimic a vegetated micro-habitat (VMH). The experiment took place over four consecutive nights and days. Groups of individuals change terrariums vertically on the second day (D2) and horizontally on the third day (D3).

Dorsal-side photographs (i.e dorsal-side reflectance) were taken at each session, from the night 1 (N1, capture of the individuals) to day 4 (D4, release of the individuals), excluding day 1 (D1). The geckos were placed in the terrariums on night 1 (N1) and not handled on day 1 (D1) to allow for acclimatisation. Body temperature and substrate temperature were taken on nights and days from day 2 (D2). Photographs and temperature measurements were always taken in the same time slot (from 12:30 pm during the day and from 11:30 pm at night). Substrate and individual temperatures were taken before each manipulation in order to eliminate any risk of heating related to the holding of the animals. On day 2 (D2), the geckos were changed of terrarium and substrate in order to study the effect of a change of micro-habitat on reflectance. Individuals in RMH were thus transferred to VMH and vice versa. In order to control for a potential effect of terrariums on chromatic variation, individuals were changed to a different terrarium but not to a different micro-habitat on day 3 (D3). In other words, individuals contained in RMH were transferred to another RMH terrarium and, similarly, individuals contained in VMH were transferred to another VMH terrarium. After being photographed one last time, the individuals were all released in the area of their capture on day 4 (D4).

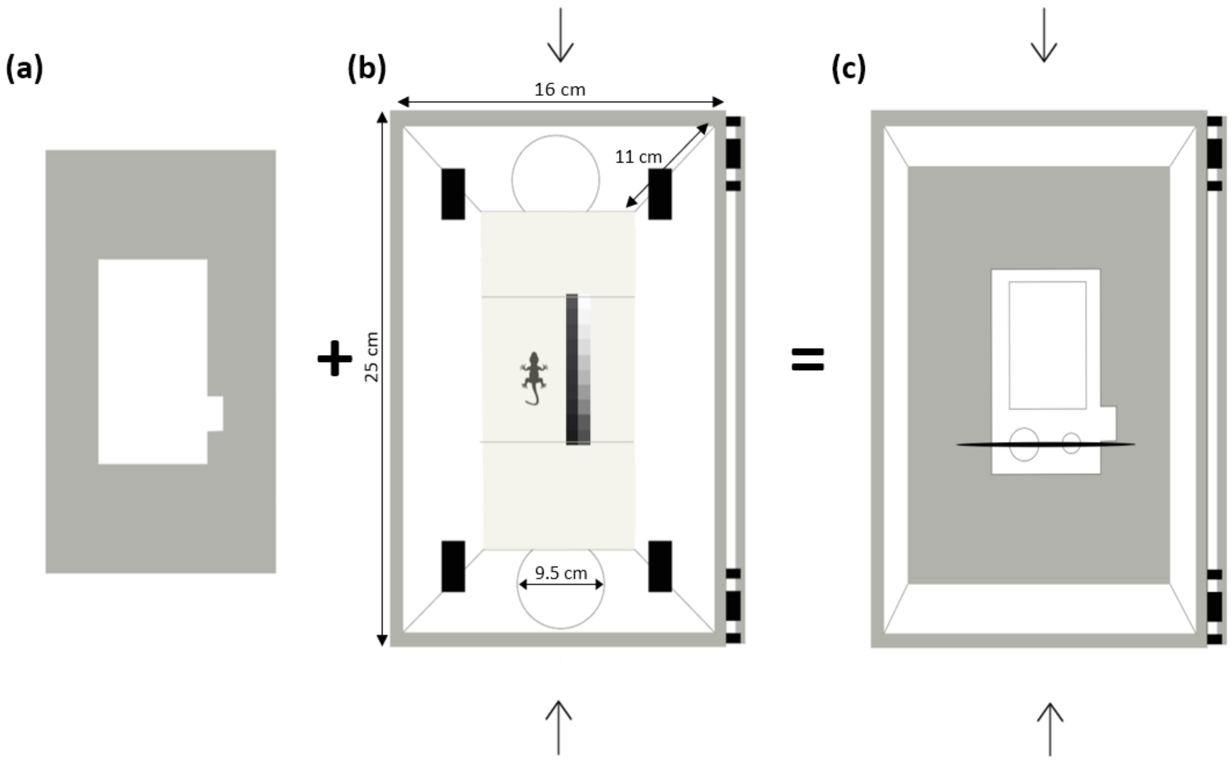


**Figure S2 :** Characteristics of the portable photographic studio (25x16x11cm) allowing the standardisation of shots. Illustration : Théo Xérès. (a) Base for the digital camera. (b) Background of the photographic studio where the individual is held next to the greyscale. The arrows indicate the holes (9.5cm diameter) for passing the hands and holding the animal. (c) Digital camera base added for shooting.

**Table S1 :** Summary of the temperature (°C) of the two substrate types, rocky micro-habitat (RMH) and vegetated micro-habitat (VMH), at night and day. For each micro-habitat, the measurements were performed on three terrariums (*N*), during two nights and three days according to the procedure specified in Figure S1. Values are given as mean ± standard deviation (SD), minimum (Min.), median and maximum (Max.).

| | **Substrate temperature (°C)** | | | | | | | | | | | |
|---|---|---|---|---|---|---|---|---|---|---|---|---|
| | **Night (*n*=2)** | | | | | | **Day (*n*=3)** | | | | | |
| **Types of substrates** | ***N*** | **Mean** | **SD** | **Min.** | **Median** | **Max.** | ***N*** | **Mean** | **SD** | **Min.** | **Median** | **Max.** |
| Rocky micro-habitat (RMH) | 3 | 16.0 | 0.7 | 15.2 | 16 | 16.8 | 3 | 24.2 | 2.8 | 21.5 | 22.7 | 29.5 |
| Vegetated micro-habitat (VMH) | 3 | 14.7 | 0.5 | 13.8 | 14.8 | 15.2 | 3 | 22.9 | 0.9 | 21.6 | 23.1 | 24.6 |

**Table S2 :** Summary of body temperature (°C) parameters of the European leaf-toed geckos at night and day. The measurements were performed on 30 individuals, during two nights and three days according to the procedure specified in Figure S1. Values are given as mean ± standard deviation (SD), minimum (Min.), median and maximum (Max.).

| | **Body temperature (°C)** | | | | | | | | | |
|---|---|---|---|---|---|---|---|---|---|---|
| | **Night (*n*=2)** | | | | | **Day (*n*=3)** | | | | |
| **Individuals** | **Mean** | **SD** | **Min.** | **Median** | **Max.** | **Mean** | **SD** | **Min.** | **Median** | **Max.** |
| 1 | 14.0 | 0.1 | 13.9 | 14.0 | 14 | 22.4 | 0.9 | 21.7 | 22.0 | 23.4 |
| 2 | 14.6 | 0.1 | 14.5 | 14.6 | 14.6 | 22.4 | 0.8 | 21.5 | 22.6 | 23.0 |
| 3 | 14.5 | 0.4 | 14.2 | 14.5 | 14.8 | 22.3 | 0.6 | 22.0 | 22.0 | 23.0 |
| 4 | 14.8 | 0.7 | 14.3 | 14.8 | 15.3 | 22.0 | 0.6 | 21.4 | 22.0 | 22.6 |
| 5 | 14.4 | 0.1 | 14.3 | 14.4 | 14.5 | 22.5 | 0.7 | 21.8 | 22.5 | 23.1 |
| 6 | 14.2 | 1.3 | 13.3 | 14.2 | 15.1 | 22.2 | 0.9 | 21.3 | 22.3 | 23.0 |
| 7 | 14.7 | 0.7 | 14.2 | 14.7 | 15.2 | 22.7 | 0.8 | 21.8 | 22.9 | 23.3 |
| 8 | 15.2 | 0.1 | 15.1 | 15.2 | 15.3 | 22.4 | 1.0 | 21.7 | 21.9 | 23.5 |
| 9 | 14.0 | 0.3 | 13.8 | 14 | 14.2 | 22.2 | 1.2 | 21.0 | 22.4 | 23.3 |
| 10 | 13.7 | 0.5 | 13.3 | 13.7 | 14.0 | 22.5 | 0.6 | 21.9 | 22.4 | 23.1 |
| 11 | 14.0 | 0.3 | 13.8 | 14.0 | 14.2 | 22.9 | 1.7 | 21.5 | 22.3 | 24.8 |
| 12 | 13.8 | 0.6 | 13.4 | 13.8 | 14.2 | 23.1 | 2.2 | 21.2 | 22.6 | 25.5 |
| 13 | 14.2 | 1.1 | 13.4 | 14.2 | 15.0 | 23.5 | 1.9 | 22.2 | 22.6 | 25.6 |
| 14 | 14.3 | 0.2 | 14.1 | 14.3 | 14.4 | 23.7 | 2.3 | 22.1 | 22.8 | 26.3 |
| 15 | 15.9 | 0.6 | 15.4 | 15.9 | 16.3 | 23.2 | 2.2 | 21.3 | 22.7 | 25.6 |
| 16 | 15.9 | 0.5 | 15.5 | 15.9 | 16.2 | 21.9 | 0.3 | 21.6 | 22.0 | 22.1 |
| 17 | 16.1 | 0.0 | 16.1 | 16.1 | 16.1 | 22.1 | 0.3 | 21.8 | 22.2 | 22.3 |
| 18 | 16.5 | 0.4 | 16.2 | 16.5 | 16.7 | 22.2 | 0.4 | 21.8 | 22.1 | 22.6 |
| 19 | 16.4 | 0.5 | 16.0 | 16.4 | 16.7 | 22.1 | 0.2 | 21.9 | 22.0 | 22.3 |
| 20 | 15.8 | 0.5 | 15.4 | 15.8 | 16.1 | 22.3 | 0.3 | 22.0 | 22.5 | 22.5 |
| 21 | 15.9 | 1.4 | 14.9 | 15.9 | 16.9 | 22.7 | 0.5 | 22.3 | 22.5 | 23.3 |
| 22 | 15.8 | 1.1 | 15.0 | 15.8 | 16.6 | 23.7 | 1.5 | 22.7 | 23.0 | 25.4 |
| 23 | 15.9 | 0.9 | 15.2 | 15.9 | 16.5 | 23.6 | 1.9 | 22.5 | 22.6 | 25.8 |
| 24 | 16.3 | 0.9 | 15.6 | 16.3 | 16.9 | 23.9 | 2.5 | 22.4 | 22.6 | 26.8 |
| 25 | 15.9 | 0.9 | 15.2 | 15.9 | 16.5 | 23.3 | 2.0 | 21.8 | 22.6 | 25.6 |
| 26 | 15.4 | 0.4 | 15.1 | 15.4 | 15.6 | 23.9 | 2.4 | 22.0 | 23.1 | 26.5 |
| 27 | 17.5 | 1.8 | 16.2 | 17.5 | 18.8 | 24.0 | 4.0 | 20.6 | 23.1 | 28.4 |
| 28 | 14.8 | 0.6 | 14.3 | 14.8 | 15.2 | 24.0 | 2.9 | 22.0 | 22.8 | 27.3 |
| 29 | 15.0 | 1.1 | 14.2 | 15.0 | 15.8 | 22.8 | 1.3 | 21.4 | 23.3 | 23.8 |
| 30 | 15.8 | 0.7 | 15.3 | 15.8 | 16.3 | 24.3 | 3.5 | 21.5 | 23.2 | 28.2 |
